# Oligodendrocyte Precursor Cells Sculpt the Visual System by Regulating Axonal Remodeling

**DOI:** 10.1101/2021.03.11.434829

**Authors:** Yan Xiao, Laura J Hoodless, Luigi Petrucco, Ruben Portugues, Tim Czopka

**Author notes:** Correspondence to, Dr. Tim Czopka, University of Edinburgh, Centre for Clinical Brain Sciences, Chancellor’s Building, 49 Little France Crescent, Edinburgh EH16 4SB, United Kingdom.

## Abstract

Many oligodendrocyte precursor cells (OPCs) do not differentiate to form myelin, suggesting additional roles of this cell population. The zebrafish optic tectum contains OPCs in regions devoid of myelin. Elimination of these OPCs impaired precise control of retinal ganglion cell axon arbor size during formation and maturation of retinotectal connectivity, and in consequence impaired visual acuity. Therefore, OPCs fine-tune neural circuit structure and function independently of their canonical role to make myelin.

## Main Text

Oligodendrocytes play crucial roles in modulating information processing through regulation of axon conduction and metabolism ^1–3^. Myelinating oligodendrocytes arise by differentiation of oligodendrocyte precursor cells (OPCs), which are uniformly distributed across the CNS and tile the tissue with their elaborate process networks ^4^. The formation of new myelin through oligodendrocyte differentiation continues into adulthood and can dynamically change to shape axonal myelination ^5–8^, However, the CNS comprises more OPCs than ever differentiate, making about 5% of all CNS cells lifelong ^9^. How this persistent population of resident CNS cells affects the CNS apart from being the cellular source of new myelin is largely unclear.

OPCs are a heterogenous population with different properties ^10,11^. Clonal analyses have shown that a large proportion of OPCs does not directly generate myelinating oligodendrocytes, suggesting that these cells may have additional physiological functions in the healthy CNS ^12^. Indeed, OPCs express molecules that can affect form and function of neurons ^13–15^, and altered gene expression in OPCs has recently been linked to mood disorders in humans ^16^. However, as changes in OPCs also affect myelination, it remains unclear if roles in the formation of a functional neural circuit can be directly attributed to OPCs that are independent of myelination.

In order to reveal myelination independent roles for OPCs, we have identified the optic tectum of larval zebrafish as a brain area which is densely interspersed with OPCs that rarely differentiate to oligodendrocytes (Fig. 1, Supplementary Fig. 1a-c, Supplementary Movies 1 and 2). Transgenic lines labeling retinal ganglion cell (RGC) axons as primary input to the tectum (Tg(isl2b:EGFP)), OPCs (Tg(olig1:memEYFP)), and myelin (Tg(mbp:memRFP)) allowed us to carry out high-resolution, whole brain imaging of neuron-oligodendrocyte interactions. Myelination of RGC axons was observed along the optic nerve and also along tectal neuron axons projecting to deeper brain areas (Supplementary Fig. 1a-c). However, the tectal neuropil where RGC axons connect to tectal neuron dendrites remained largely devoid of myelin until at least 14 days post fertilization (dpf) despite being interspersed with OPC processes throughout (Fig. 1b, c). Quantification of oligodendrocyte numbers confirmed that no more than 6% of oligodendrocyte lineage cells were differentiated by 14 dpf, which was in strong contrast to hindbrain regions showing 56% differentiation (Supplementary Fig. 1d, e). Within the tectum, OPCs localized their soma either at the border between tectal neuropil and periventricular zone containing the majority of tectal neurons or right within the neuropil (Fig. 1d-f, Supplementary Fig. 1f), and claimed non-overlapping territories, similar to previous studies (Supplementary Fig. 1g, Supplementary Movie 3) ^12,17,18^. Being a non-differentiated cell type, OPCs can be highly dynamic and potentially migrate, proliferate, or differentiate. However, sparse labeling revealed that soma positioning of individual tectal OPCs remained largely stable with a low rate of division and differentiation between 6-10 dpf (Fig. 1f-g).

**Fig. 1.**
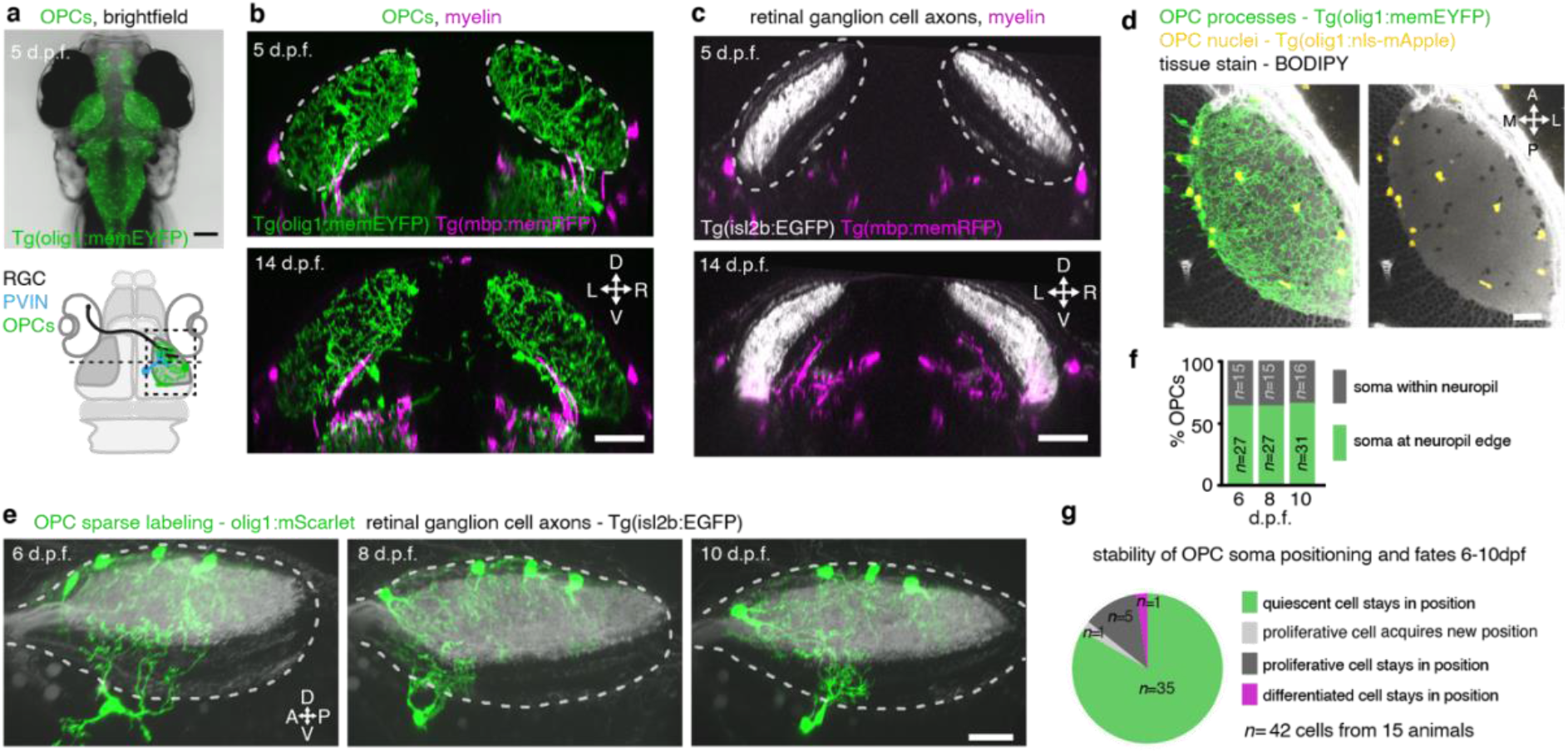
The tectal neuropil of larval zebrafish is interspersed with OPC processes but largely devoid of myelin. **a)** Transgenic zebrafish line showing OPC processes throughout the brain. Dashed line indicates cross-sectional plane shown in b and c. Schematic of larval zebrafish brain delineates RGC axons, dendrites of periventricular interneurons (PVINs), and OPCs. **b** and **c)** Cross sectional views of transgenic zebrafish showing that OPC processes intersperse the tectal neuropil (dashed outlines) while myelin is largely absent. Scale bars: 50 μm. **d)** Subprojection of OPC reporter lines (dorsal view) stained with BODIPY to outline tectal neuropil and nuclear regions. Scale bar: 20 μm. **e)** Time-lapse of four individual OPCs (lateral rotation view) in transgenic RGC reporter animal. Dashed outlines depict tectal neuropil. Scale bar: 20 μm. **f** and **g**) Quantifications of individual OPCs as shown in e showing low rates of soma position changes, division and differentiation.

The appearance of OPCs coincided with the time when RGC axons arrive in the developing tectum and establish their terminal arborizations (Supplementary Fig. 2). Time-lapse imaging revealed that RGC arbors dynamically interact with OPC processes with continuous contacts and repulsions (Fig. 2a, Supplementary Movie 4). This spatial and temporal correlation prompted us to ask whether OPCs influence the organisation of RGC arbors within the tectal neuropil. To test this, we used an inducible Nitroreductase (NTR)-mediated cell ablation system specifically targeted to OPCs (Tg(olig1:CFP-NTR)) (Fig. 2b, c, Supplementary Fig. 3a). Early OPC ablation from 2dpf, the time when RGC axons arrive at the tectum, led to formation of erroneous axon branches reaching outside the tectal neuropil, as well as enlarged arbor sizes of individual RGC axons (Fig. 2e, f, h, i). To exclude that this phenotype was mediated indirectly by microglia clearing dying OPCs, or by diverting microglial activities which have an established role at eliminating synapses ^19^, we carried out two controls: genetic depletion OPC numbers without causing inflammation by morpholino injection against *olig2*, and depletion of microglia using a morpholino against *interferon regulatory factor 8* (*irf8*) (Supplementary Fig. 3b-d). While *olig2* morphants also exhibited ectopic branching and enlarged RGC arbors, none of these phenotypes were seen *irf8* morphants lacking microglia (Fig. 2g, j). Therefore, erroneous RGC arborizations and RGC arbor size resulted directly from the absence of OPCs.

**Fig. 2.**
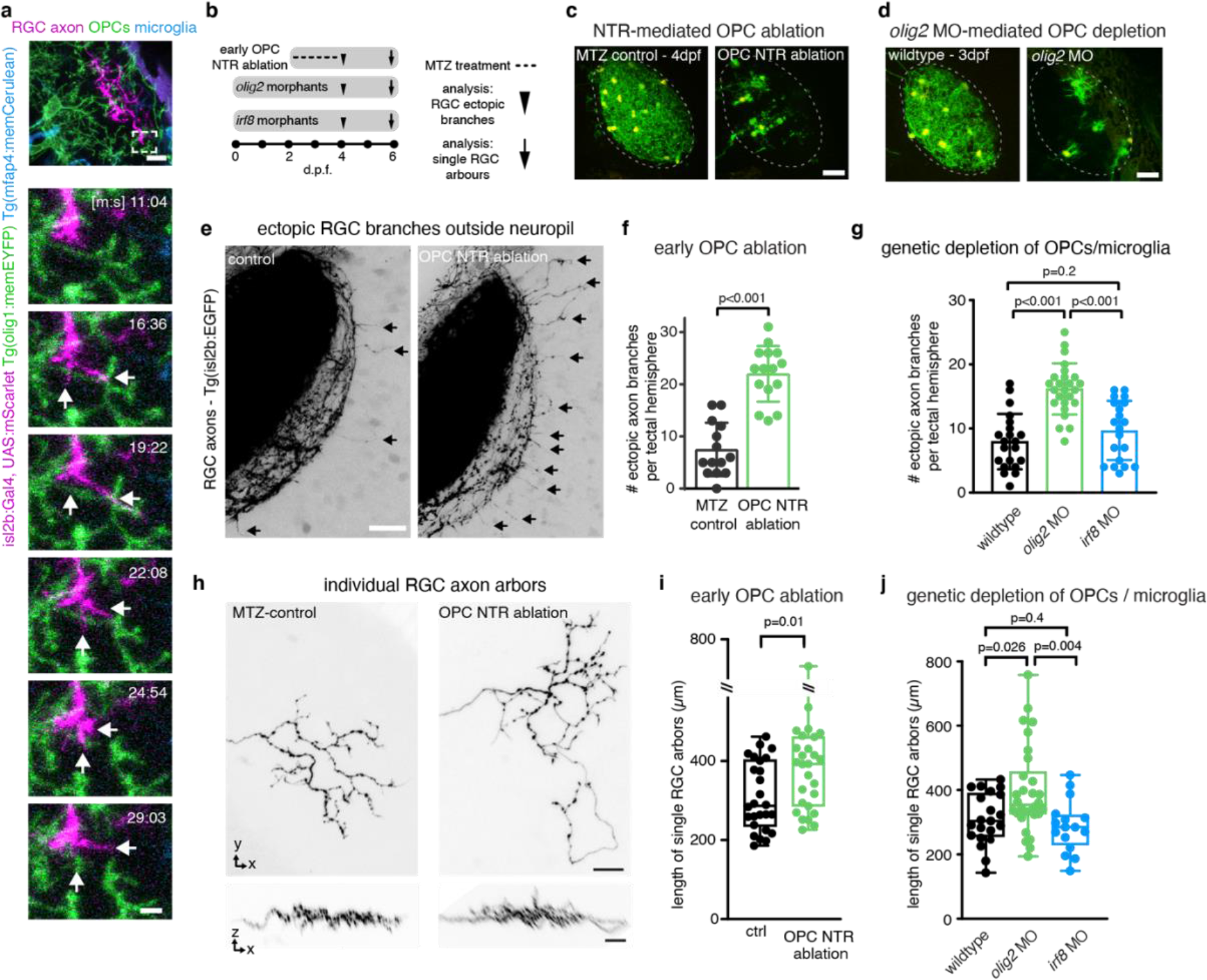
Early OPC depletion causes formation of aberrant RGC arborizations. **a)** Time-lapse showing dynamic contacts and retractions between RGC axon arbors and OPC processes (arrows). Scale bars: 10 μm (top), 2μm (bottom). **b)** Timelines of manipulations in this figure. **c** and **d)** Example images of NTR-mediated OPC ablation (c) and olig2 morpholino-mediated OPC depletion. Dashed lines outline tectal neuropil. Scale bars: 25μm. **e** and **f)** Increased formation of ectopic RGC axon branches extending outside tectal neuropil (arrows) upon early ablation of OPCs (mean 7.5±5.1 S.D. in control *vs*. 22.0±5.3 in OPC-NTR ablation, n=14/15 animals, unpaired two-tailed t test). Scale bar: 20 μm. **g)** Increased formation of ectopic RGC axon branches upon genetic OPC reduction (*olig2* morphants) but not upon microglial depletion (*irf8* morphants) (mean 7.9±4.3 S.D. in control *vs*. 16.2±3.9 in *olig2* MO *vs*. 9.7±4.6 in *irf8* MO, n=21/25/20 animals, one-way ANOVA). **h** and **i)** Increased size of single RGC arbors upon early OPC ablation (top) whilst maintaining single lamina layering (bottom) (median 287±403/235 I.Q.R. in control *vs*. 393±461/286 in OPC-NTR ablation, n=26/27 axons in 24/23 animals, two-tailed Mann-Whitney test). Scale bars: 10 μm. **j)** Increased size of single RGC arbors upon genetic OPC depletion (*olig2* morphants) but not upon microglial depletion (*irf8* morphants) (median 304±391/255 I.Q.R. in control *vs*. 355±458/326 in *olig2* MO *vs*. 285±323/229 in *irf8* MO, n=21/30/16 axons in 11/14/12 animals, Kruskal Wallis test)

Following their formation, RGC arbors undergo a phase of developmental pruning during larval stages when retinotectal connectivity is refined ^20,21^. To test whether OPCs also play a long-lasting role during the refinement of RGC arbors as they continue to persist and interact with each other, we carried out late OPC ablations starting from 7dpf when zebrafish have a functional visual system. Ablations were carried out analogously using NTR-mediated chemogenetics, or by 2- photon mediated cell ablation to specifically eliminate OPCs from the tectum (Fig, 3, Supplementary Fig. 4). In control animals, individual RGC arbors underwent process remodelling with additions and eliminations of multiple neurites that lead to a net reduction in arbor size by about 14% between 7 and 10 d.p.f., similar to previous reports (Fig. 3c, Supplementary Fig. 4g, 5a-c) ^21^. This reduction in arbor size was significantly decreased in OPC-ablated animals (1.7 % after OPC laser ablation, 6% after OPC NTR ablation), with some arbors even increasing in size due to a reduction in neurite eliminations as well as an increase in neurite additions (Fig. 3c, Supplementary Fig. 4g, 5a-c). Despite changes in size, individual arbors maintained restricted to single tectal laminae upon OPC depletion (Supplementary Fig. 4g, 5b). This is distinct from the role of radial glia which are required to maintain stratification of the tectal neuropil ^22^. Furthermore, the effects on neurite remodelling were specific to axonal processes within the tectum because remodelling of tectal neuron dendrites in the same tissue was unaffected upon OPC ablation, further corroborating that the effects observed on RGC arbors do not result from unspecific collateral damage induced by our manipulations (Supplementary Fig. 5d-g).

**Fig. 3.**
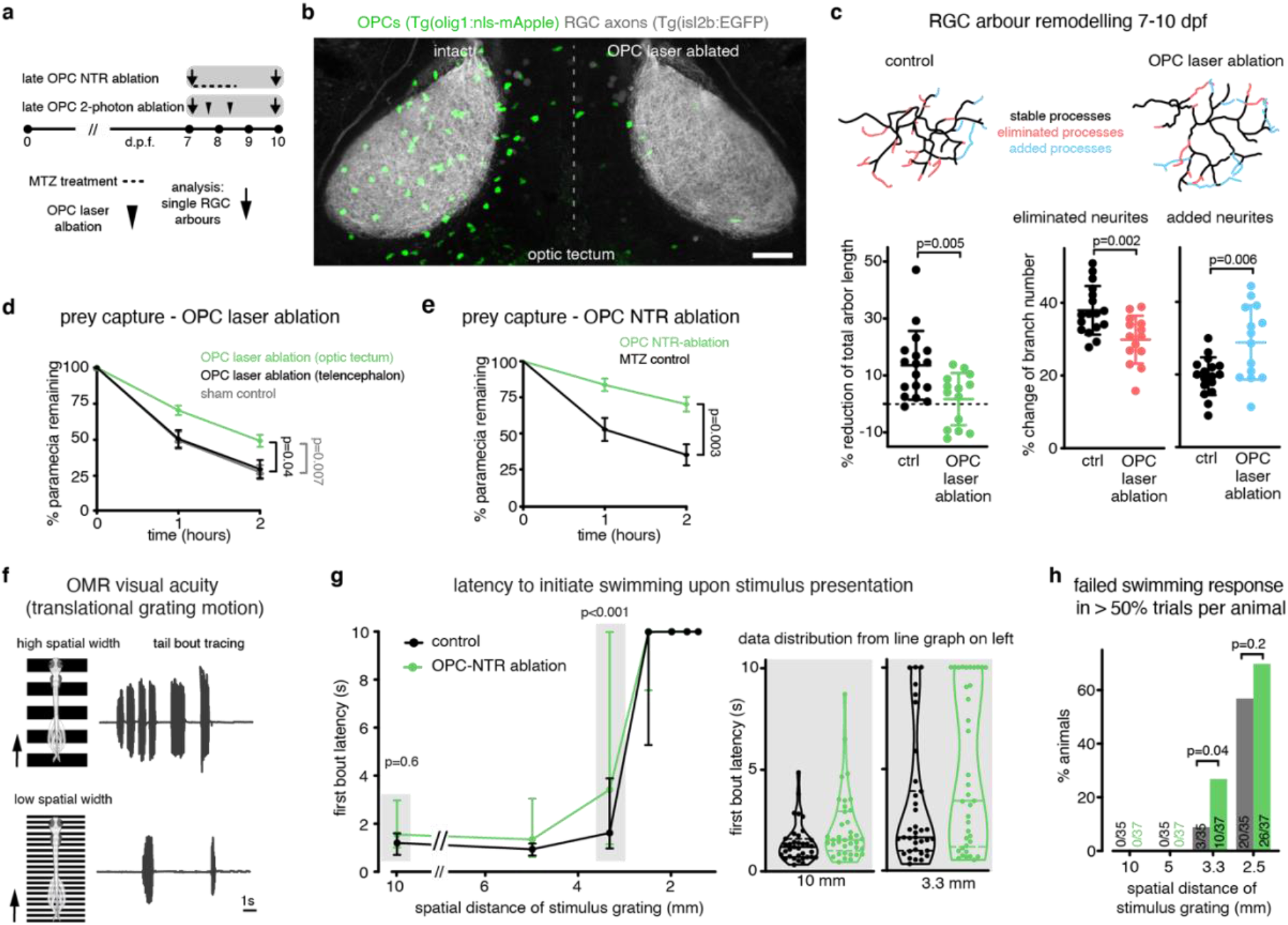
Late OPC ablation impairs RGC arbor remodeling, prey capture behavior and visual acuity. **a)** Timelines of manipulations for late OPC ablations. **b)** Dorsal views showing unilateral laser ablation of OPCs in the tectum. Scale bar: 20 μm. **c)** Reconstructions of time-projected RGC arbors highlighting stable, eliminated, and added processes. Quantifications show diminished developmental reduction of RGC arbors between 7 and 10 dpf in OPC-ablated animals (left graph), mediated by decreased branch eliminations (middle) and enhanced additions (right) (left: mean 13.6±12.1 S.D. in control *vs*. 1.7±9.2 in OPC laser ablation; middle: 37.9±6.7 *vs*. 29.9±6.6; right: 19.5±5.2 *vs*. 28.8±10.2; n=17/14 cells in 11/9 animals (unpaired two-tailed t test). **d** and **e)** Impaired paramecium capture rates upon tectal OPC laser ablation (d) and OPC NTR ablation (e); (d: mean 27.6±4.9 S.E.M. in sham control *vs*. 29.7±6.3 in telencephalic OPC ablation *vs*. 49.2±4.2 in tectal OPC ablation at two hour time point, n=11/10/23 animals, two-way ANOVA); (e: mean 35.2±7.3 SEM in MTZ control *vs*. 70.2±5.1 in OPC NTR ablation at two hours time point, n=18 animals per group, two-way ANOVA). **f)** Experimental setup of OMR elicited by moving gratings of different spatial widths and example trace of tail bout recording. **g)** First bout latencies in OMR assays. Violin plots show distribution of individual data points at 10 mm and 3.3 mm spatial frequency (10mm: median 1.2±1.6/0.7 I.Q.R. in control *vs*. 1.5±2.9/1.0 in OPC-NTR ablation; 3.3mm: median 1.6±3.9/0.9 in control *vs*. 3.4±9.9/1.1 in OPC-NTR ablation; n=35/37 animals, two-way ANOVA) **h)** Enhanced failure to initiate swimming in response to narrow moving gratings upon OPC NTR ablation (one-tailed Fisher’s exact test).

It has been reported that enlarged RGC arbor sizes reduce visual acuity and in consequence impair the ability to capture prey ^23^, which is a visually evoked behaviour that requires RGCs projections into superficial tectal layers ^24,25^. To test if OPC-mediated changes in RGC axon growth are of relevance for visual processing, we assayed prey capture behaviour and visual acuity. Late OPC ablation significantly reduced prey capture success rates, with about two times more paramecia remaining after NTR ablation, and about 70% more paramecia remaining uncaptured after unilateral laser ablation of tectal OPCs (Fig. 3d, e, Supplementary Fig. 6a, b). This effect was specific to tectal OPCs as OPC laser ablation from adjacent telencephalic regions did not impair paramecium capture (Fig. 2d, Supplementary Fig. 6a). None of our manipulations affected overall swimming activity, ruling out that gross locomotor defects account for reduced hunting (Supplementary Fig. 6c, d). To further test if OPC ablation impairs visual processing, we performed optomotor response (OMR) assays stimulated by moving gratings of different spatial frequencies (Supplementary Fig. 6e, f, Supplementary Movie 5). Higher spatial frequencies become increasingly difficult to resolve, leading to longer latencies and ultimate failure to elicit an OMR (Fig. 3f). We found that OPC ablated animals were able to robustly elicit OMR at lower spatial frequencies, but increasingly failed to show a swimming response when high spatial frequency stimuli were presented (Fig. 3g, h, Supplementary Fig 6g-j). The enhanced failure to respond to visual stimuli that are increasingly difficult to distinguish indicate that OPC-ablated zebrafish have impaired visual acuity.

Together, our data reveal a role for OPCs in fine-tuning the structure and function of neural circuits that is independent of their traditional role in myelin formation, and which is mediated by regulating growth and remodeling of axon arbors. This provides physiological function for OPCs cells that is beyond their role simply being a cellular source for myelinating oligodendrocytes, or inhibiting regeneration in glial scars, but rather complements the roles of microglia and astrocytes in shaping neuronal form and function. Future work will show if this non-canonical role of resident OPCs is a feature that is shared by all OPCs or if it is rather restricted to subsets of cells, areas of the CNS, and mechanisms of action.

## Acknowledgments

We are grateful to Wenke Barkey for excellent technical assistance with molecular cloning and zebrafish husbandry, and to Thomas Misgeld for providing generous access to 2-photon microscopy equipment. We thank Rafael Almeida, David Lyons, and all members of the Czopka lab for critical input and discussion of the manuscript. We thank Martin Meyer (King’s College London) for the foxp2A promoter plasmid, David Lyons (University of Edinburgh) for the olig2 morpholino, and Leanne Godinho/Rachel Wong (TU Munich/Washington University) for the isl2b:Gal4 plasmid. This work was funded by an ERC Starting grant (MecMy #714440) to TC, the German Research Foundation (DFG) SFB870/TP A14 and the Emmy Noether Programme (Cz226/1-1 and Cz226/1-2) to TC, the DFG under Germany’s Excellence Strategy within the framework of the Munich Cluster for Systems Neurology (EXC 2145 SyNergy #390857198) to TC and RP, and the TRR 274/1 2020 (#408885537) to RP.

## Author contributions

Conceptualization: TC; Methodology: YX, LP, LJH; Software: LP and RP; Investigation: YX and LJH; Formal analysis: YX, LP, LJH; Supervision: RP and TC; Visualization: YX and TC; Writing - original draft: YX and TC; Funding acquisition: RP and TC.

## Material and Methods

### Zebrafish lines and husbandry

We used the following existing zebrafish lines and strains: Tg(mbp:nls-EGFP)^zf3078tg 1^, Tg(mbp:memRFP)^tum101tg 2^, Tg(mbp:memCerulean)^tum102tg 2^, Tg(olig1:memEYFP)^tum107tg 3^, Tg(olig1:nls-mApple)^tum109tg 3^, Tg(olig1:nls-Cerulean)^tum108tg 3^, Tg(mfap4:memCerulean)^tum104tg 3^, Tg(isl2b:EGFP)^zc7tg 4^, AB and nacre. The transgenic line Tg(olig1:CFP-NTR) was newly generated for this study. All animals were kept at 28.5 °C with a 14h-10h light-dark cycle according to local animal welfare regulations. All experiments carried out with zebrafish at protected stages have been approved by the government of Upper Bavaria (animal protocols ROB-55.2-1-54-2532.Vet_02-18- 153, ROB-55.2-2532.Vet_02-15-199, and ROB-55.2-2532.Vet_02-15-200 to T.C.).

### Transgenesis constructs

To generate the middle entry clone pME_EYFP, the coding sequence was PCR amplified from a template plasmid using the primers attB1_YFP_F: GGGGACAAGTTTGTACAAAAAAGCAGGCTGCCACCATGCTGTGCTGC and attB2R_YFP_R: GGGGACCACTTTGTACAAGAAAGCTGGGTCTTACTTGTACAGCTCGTCCATGC.

The PCR product was recombination cloned into pDONR221 using BP clonase (Invitrogen).The expression constructs pTol2_olig1:mScarlet, pTol2_olig1:EYFP, pTol2_olig1:tagCFP, pTol2_olig1:tagCFP-NTR, pTol2_cntn1b:mScarlet, and pTol2_foxp2A:mScarlet were generated in multisite LR recombination reactions with the entry clone described above, p5E_olig1 ^2^, p5E_cntn1b ^5^, p5E_foxp2A (gift from Dr. Meyer M.P., King’s College London) ^6^, pME_tagCFP ^2^, pME_mScarlet ^3^, p3E_NTR-pA ^1^, p3E_pA and pDestTol2_pA of the Tol2Kit ^7^. The expression construct pTol2_isl2b:Gal4 using a published isl2b promoter clone ^8^ was a kind gift of Dr. Leanne Godinho (TU Munich) and originally provided by Dr. Rachel Wong (Washington University); pTol2_olig1:memEYFP ^3^, and pTol2_10xUAS:mScarlet ^3^ have been published previously.

### DNA microinjection for sparse labeling and generation of transgenic lines

Fertilized eggs at one-cell stage were microinjected with 1 nl of a solution containing 5-20 ng/μl DNA plasmid and 20 ng/μl Tol2 transposase mRNA. Injected F0 animals were either used for single-cell analysis or raised to adulthood to generate full transgenic lines. For this, adult F0 animals were outcrossed with wild-type zebrafish, and F1 offspring were screened for presence of the reporter transgene under a fluorescence stereo dissecting microscope (Nikon SMZ18).

“Tg(promoter:reporter)” denotes a stable transgenic line, whereas “promoter:reporter” alone indicates that a respective plasmid DNA was injected for sparse labeling of individual cells.

### OPC ablation using Nitroreductase (NTR)

For NTR mediated OPC ablation at early developmental stages, Tg(olig1:CFP-NTR) zebrafish at 2dpf were incubated in 10mM metronidazole (MTZ) dissolved with 0.2% DMSO in 0.3× Danieau’s solution for 48h at 28°C in the dark, with a change of solution after 24h. After MTZ incubation, embryos were rinsed and kept in 0.3×Danieau’s solution until analysis.

For NTR-mediated OPC ablation at later larval stages, Tg(olig1:CFP-NTR) zebrafish at 7dpf were incubated in 10mM metronidazole (MTZ) dissolved with 0.2% DMSO in 0.3× Danieau’s solution for 24h at 28°C in the dark. After MTZ incubation, larvae were rinsed and kept in nursery tanks with standard diet until 10 dpf. Non NTR-expressing zebrafish treated with 10 mM MTZ were used as controls in all experiments.

### OPC ablation using two-photon lasers

OPCs were laser ablated from Tg(olig1:nls-mApple) using an Olympus FV1000/MPE equipped with a MaiTai DeepSee HP (Newport/Spectra Physics) and a 25x 1.05 NA MP (XLPLN25XWMP) water immersion objective. Continuous confocal scans using a 559nm laser were taken to locate individual OPC nuclei in the optic tectum, which was identified by additional transgenic genetic markers (Tg(olig1:memYFP) or Tg(isl2b:EGFP)). Each cell was ablated using a 500 ms line scan across the cell nucleus using the MaiTai laser tuned to 770nm (1.75W output). The wavelength was incrementally increased for ablating OPCs in deeper tissue. After successful ablation, previously bright, round nuclei appeared dim, irregular or fragmented. The ablation procedure was repeated when cells did not show this signature. Unilateral OPC ablations took 60-90min in the tectum. For analyzis of axon remodeling after OPC laser ablation, surviving and/or repopulating OPCs were ablated again on the second day.

### Morpholino-mediated depletion of microglia and OPCs

Microglia were depleted from zebrafish embryos by microinjection of 4.5pg of a previously published morpholino targeting the start codon of *irf8* (5’-TCAGTCTGCGACCGCCCGAGTTCAT-3’) ^9^. OPCs were depleted by microinjection of 7.5pg of a previously published morpholino targeting the start codon of *olig2* (5’-ACACTCGGCTCGTGTCAGAGTCCAT-3’) ^10^. Both morpholinos were synthesized by Gene Tools.

### Neutral red staining

Zebrafish embryos were incubated for 2.5 h in the dark in 2.5μg/ml neutral red solution (Sigma Aldrich, n2889) diluted in Danieau’s solution. Afterwards, embryos were washed 3 times 10 minutes with Danieau’s solution. Brightfield images of the head of the fish were taken using a Leica DFC300 FX Digital Color Camera.

### Prey capture assay

2 ml 0.3×Danieau’s solution with 30 Paramecium multimicronucleatum were added to a 35mm dish, along with a single zebrafish larva. The number of remaining paramecia was determined at hourly intervals for 2 hours. To rule out batch dependent effects resulting from “natural” paramecia death followed by their disintegration, a control containing paramecia but no fish was run alongside each experiment. Spontaneous paramecia death occurred only rarely in 0-3%.

### Locomotor activity assay

For all experiments, testing occurred between 9 am and 5 pm using a randomized trial design to eliminate systematic effects due to the time of day. A tracking chamber was prepared by using a 35 mm petri dish mold surrounded by 1% agarose situated in the center of a 85 mm petri plate to eliminate mirroring that occurs at the wall of a plastic petri dish. Single zebrafish were placed into the well filled with 2 ml 0.3× Danieau’s solution. The plate was positioned above an LED light stage to maximize contrast for facilitating zebrafish tracking (two dark eyes and swim bladder of zebrafish larva on a light background), with a high-speed camera (XIMEA MQ013MG-ON) equipped with a Kowa LM35JC10M objective positioned above the dish. The larvae acclimated to the recording arena for 5 min before the start of video acquisition. The center of mass of two eyes and swim bladder was taken as the center of the fish. Subsequently, video of spontaneous freely-swimming was recorded for 10 min at 100 Hz using a custom written Python script and the Stytra package ^11^.

### Optomotor response assay

Zebrafish larvae were embedded in 1% agarose in a 35 mm petri dish. After allowing the agarose to set, the dish was filled with 0.3× Danieau’s solution and the agarose around the tail was removed with a scalpel, leaving the tail of fish free to move (hereby referred to as a head-restrained preparation) {Portugues:2011in}. Visual stimuli were presented on the screen from below using an Asus P3E micro projector and an infrared light (Osram 850 nm high power LED). The fish’s tail was tracked using a high-speed camera (XIMEA MQ013MG-ON) and a 50mm telecentric objective (Navitar TC.5028). A square-wave grating with variable spatial period and maximal contrast achieved by the projector (black and white bars), and online tail tracking and stimulus control was performed using Stytra software 0.8.26 ^11^. Experiments were performed in closed-loop, meaning that the behavior of fish was fed back to the visual stimulus to provide the fish with visual feedback. Therefore, when the fish swam, the grating accelerated backward at a rate proportional to swim power, i.e., [stimulus velocity] = 10 – [gain] × [swim power]. Swim power was defined as the standard deviation of the tail oscillation in a rolling window of 50 ms. To obtain a feedback that mimics the visual feedback the animal would receive when freely swimming, the gain multiplication factor was chosen to result in an average fictive velocity of about 25 mm/s during the bout. When the fish was not swimming [swim power] = 0, the grating moved in a caudal to rostral (forward) direction at a baseline speed of 10 mm/s. The stimulus scene was a square window that was centered on the head of fish and spanned a field of total 60 × 60 mm. For analysis, individual bouts were counted as episodes where the swim power was above 0.1 rad for at least 100 ms. Then latency to first bout and total number of bouts were quantified for each trial (Latency was set a default value equal to the stimulus duration, when fish did not respond). Analysis was performed with custom scripts written in Python.

### Zebrafish mounting for live-cell microscopy

Zebrafish larvae were anaesthetized with 0.2 mg/ml 3-aminobenzoic acid ethyl ester (MS-222). For confocal microscopy, animals were mounted ventral side up in 1% ultrapure low melting point agarose (Invitrogen) onto a glass-bottom 3-cm Petri dish (MatTek). For two-photon microscopy, embryos were mounted ventral side up in low melting point agarose on a glass coverslip. The coverslip was then flipped over on a glass slide with a ring of high-vacuum grease filled with a drop of 0.2 mg/ml MS-222 to prevent drying out of the agarose. After imaging, the animals were either euthanized or released from the agarose using microsurgery forceps and kept individually until further use.

### Immunohistochemistry

Samples were fixed overnight at 4°C in a solution of 4% paraformaldehyde in phosphate-buffered saline (PBS) solution containing 1% Tween-20 (PBST). After fixation, the samples were washed in the same solution without fixative and blocked for 1.5 h at room temperature in PBS buffer, 0.1% Tween20, 10% FCS, 0.1% BSA and 3% normal goat serum. Primary antibody incubation was conducted at 4°C overnight in blocking solution. Afterwards, samples were washed three times in PBS with 0.1% Tween20 and then incubated with Alexa Fluor 633-conjugated secondary antibody. Stained samples were washed three times in PBS with 0.1% Tween20, and subsequently mounted with ProLong Diamond Antifade Mountant (Thermo Fisher Scientific). Primary antibody rabbit anti-HuC/D (abcam ab210554) were used at a dilution of 1/100. Goat anti-rabbit secondary antibody (Thermo Fisher) were used at a dilution of 1:1000. Images were obtained using a confocal microscope (Leica TCS SP8).

### Confocal microscopy

Images of embedded zebrafish were taken at a Leica TCS SP8 confocal laser scanning microscope. We used 458 nm for excitation of Cerulean and tagCFP; 458 and 488 nm for EGFP; 514 nm for EYFP; 561 nm for mApple and mScarlet; 633 nm for AF633 and BODIPY630/650. For overview images and analysis of cell numbers based on nuclear transgenes, we used a 10× /0.4 NA objective (acquisition with 568 nm pixel size (xy) and 2 μm z-spacing). For all other analyses, we acquired 8 or 12 bit confocal images using a 25× /0.95 NA H2O objective with 114-151 nm pixel size (xy) and 1 or 1.5 μm z-spacing.

### Analysis of axonal and dendritic arbor remodeling

Only RGC axons that arborized within the tectal neuropil and which could be traced back to the optic nerve were included for analysis. The neurites of PVIN in the superficial layer were randomly selected for analysis, as the dendrites of PVIN located in the superficial layer and their axons located in deeper layer {Robles:2011}. Individual axonal and dendritic arbors were analyzed using the segmentation tool of 3D tracing in the simple neurite tracer plugin in Fiji/ImageJ ^12^. Each arbor was traced from its first branch point out to all branch tips, and each branch segment was counted from the branch point to the next branch point or branch tips. From this tracing, we extracted the measurement of total branch length and the total number of branch segments. The eliminated/added branch segment was obtained by comparing the tracing of the same neuron at two time points.

### Image and data presentation

Images were analyzed with Fiji and Imaris. Morphology reconstructions were carried out with the Imaris FilamentTracer module. Data were prepared and assembled using Graphpad Prism 7 and 8, Fiji, and Adobe Illustrator CS6 and 2020.

### Statistics and reproducibility

For analyses that involved cohorts of animals or treatment groups, zebrafish embryos of all conditions were derived from the same clutch and selected at random before treatment. No additional randomization was used during data collection. For time-course analyses of OPCs, zebrafish were screened for single-cell labeling before imaging, and all animals with appropriate expression were used in the experiment. No data were excluded from the analyses. We selected sample sizes based on similar sample sizes that have previously reported ^13–16^. No statistical analysis was used to pre-determine sample sizes. Analysis was performed using Microsoft Excel and GraphPad Prism. All data were tested for normal distribution using the Shapiro–Wilk normality test before statistical testing. In the figures, bar graphs are shown as mean ± standard deviation (S.D.); line plots are shown as mean ± SEM, or median ± interquartile range (I.Q.R.); box and whiskers plots are expressed as median ± I.Q.R., minimum and maximum values; violin plots represent the median ± I.Q.R., minimum and maximum values. For statistical tests of normally distributed data that compared two groups, we used unpaired t-tests. Non-normally distributed data were tested for statistical significance using the Mann-Whitney U test (unpaired data). For multiple comparisons test, one-way analysis of variance (ANOVA) was used for parametric data and the Kruskal-Wallis test for non-parametric data (both with Benjamini-Krieger and Yekutieli correction). Repeated measurements were tested using two-way ANOVA with Sidak’s corrections. We used Fisher’s exact test to analyze contingency tables.

---

**Supplementary Figure 1:**
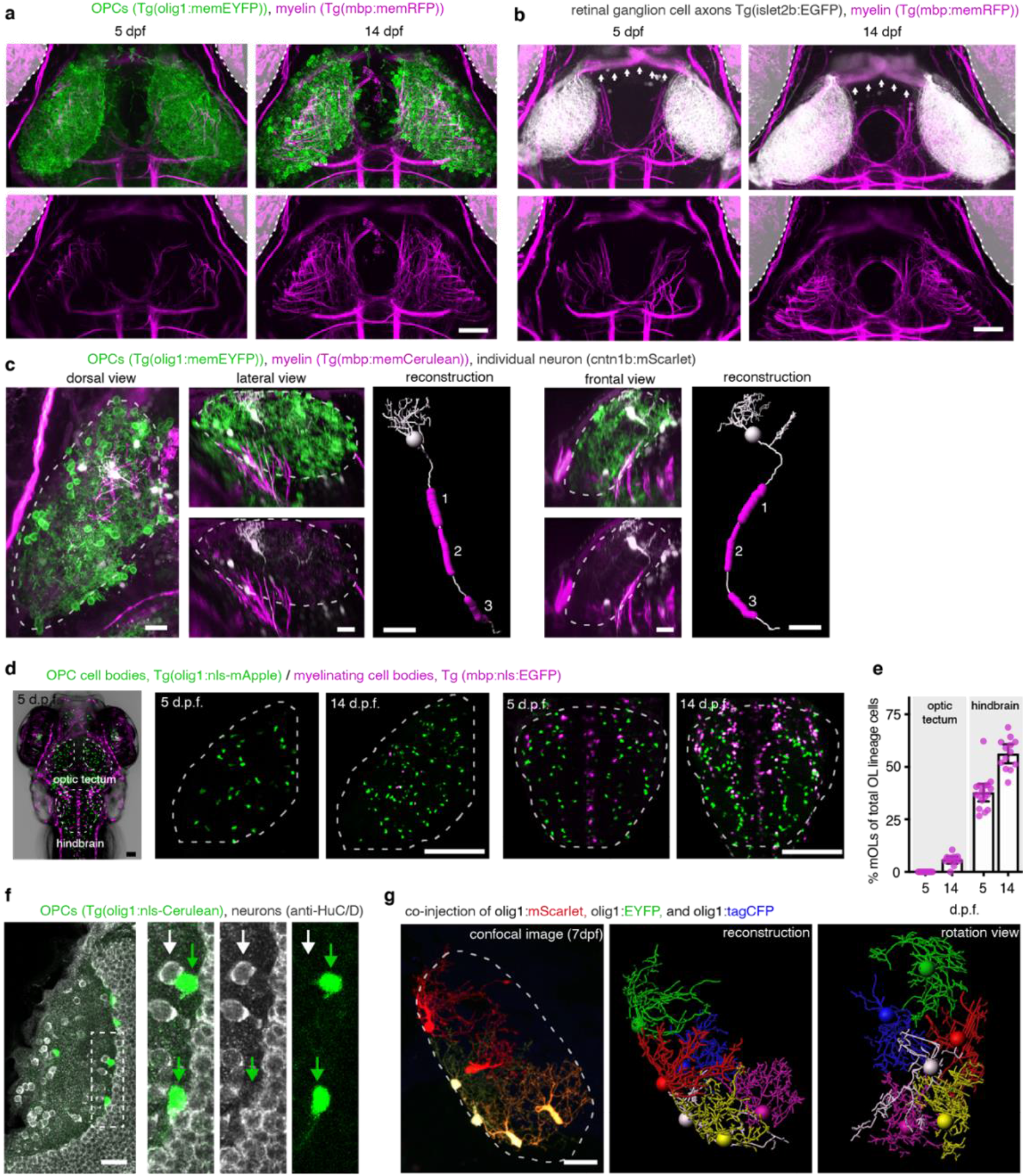
Characterization of OPCs and myelin in the zebrafish tectum. **a** and **b)** Confocal images showing dorsal views of the zebrafish visual system at 5 and 14 dpf in transgenic reporter lines for OPCs, myelin, and RGC axons as indicated. Dashed lines and grayed areas indicate position of the eyes. Arrows in (b) point to myelinated optic nerves. Scale bars: 50 μm. **c)** Confocal images and 3D reconstructions of individual neuron soma residing within the tectal neuropil and extending intermittently myelinated axon out of the tectal neuropil into deeper brain regions at 9 dpf. Dashed lines outline the tectal neuropil. Scale bars: 20 μm. **d** and **e)** Transgenic reporter animals showing the ratio between mOLs and OPCs in the optic tectum and hindbrain (n=14/12 at 5/14 dpf in tectum, and n=17/13 at 5/14 dpf in hindbrain). Scale bars: 50 μm. **f)** Whole mount immunohistochemistry showing that OPCs labeled in transgenic olig1 reporter lines (green arrows) are negative for the pan-neuronal maker HuC/D (white arrows). Scale bar: 20 μm. **g)** Multicolor labeling and 3D reconstruction of OPCs in the tectal neuropil showing that processes of individual cells occupy non-overlapping territories at 7 dpf. Dashed lines outline the tectal neuropil. Scale bar: 20 μm

**Supplementary Figure 2:**
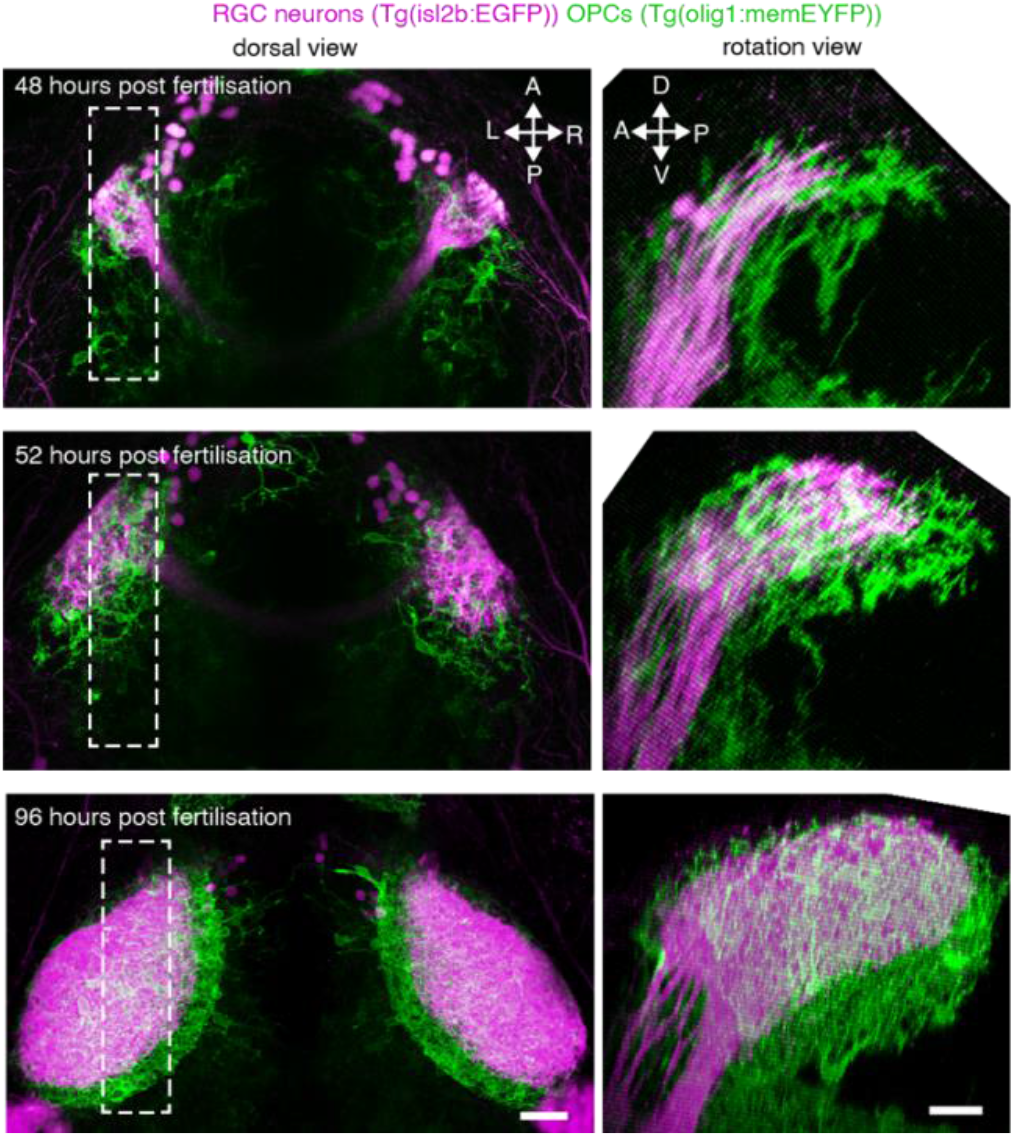
Presence of OPCs during tectal development. Dorsal (left) and rotation (right) views of confocal images showing the presence of OPCs from 48 hours post fertilization when RGC axons arrive in the tectum. Dashed rectangles indicate area shown in rotation views. Scale bars: 20 μm.

**Supplementary Figure 3:**
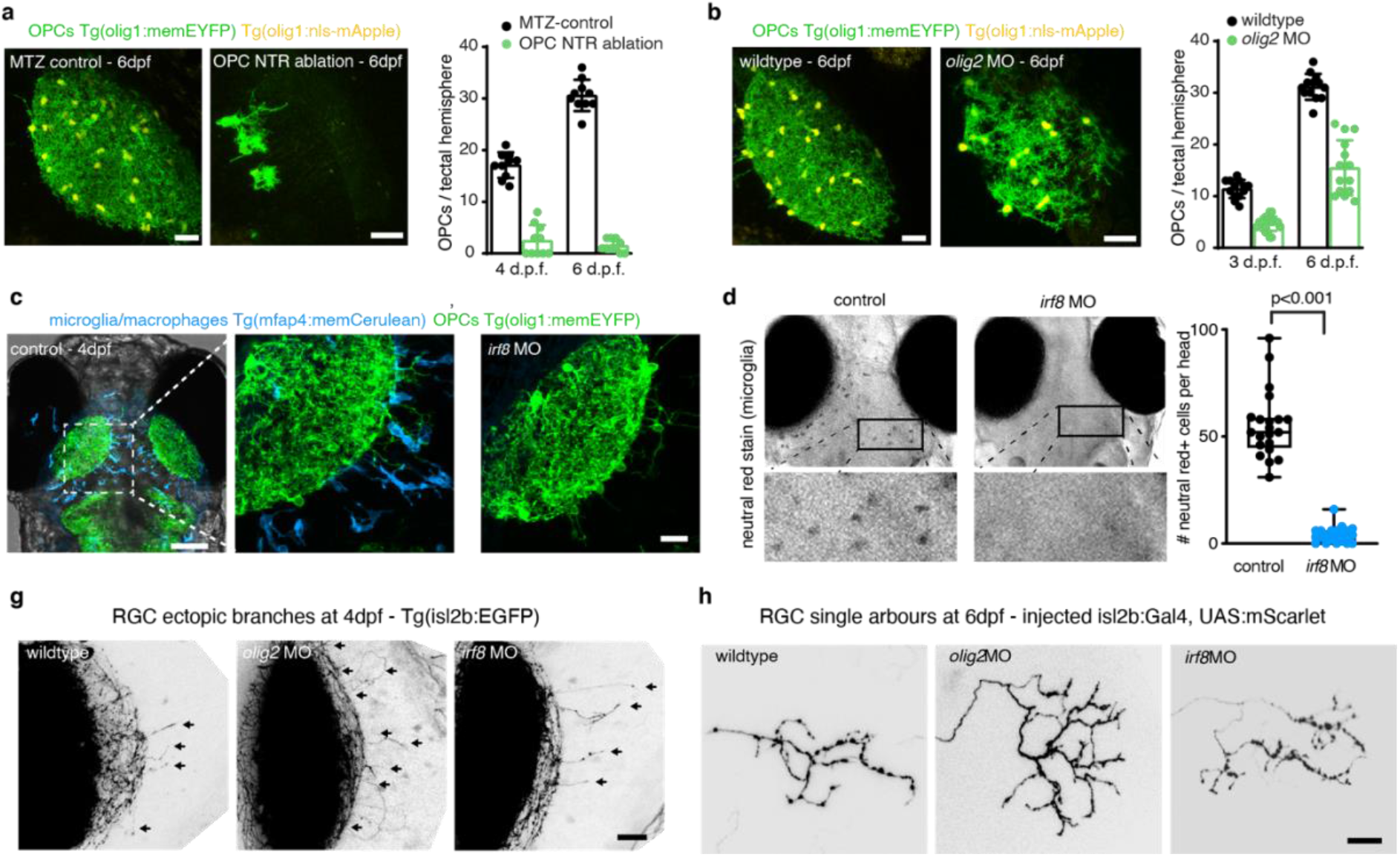
Early ablation and depletion of OPCs and microglia. **a)** Example images of OPCs in tectal hemisphere and quantifications revealing efficient ablation from in olig1:CFP-NTR animals upon MTZ application from 2dpf (n=10 animals per group). Scale bar: 25 μm. **b)** Example images of OPCs in tectal hemisphere and quantifications revealing genetic depletion of OPCs in *olig2* morphants (n=13/14 animals in wildtype/*olig2* MO). Scale bar: 25 μm. **c)** Dorsal views of transgenic reporter lines showing presence of OPC within the tectal neuropil while microglia largely reside outside the neuropil. Dashed box indicates area shown in middle panel. Scale bar: 100 μm. Right panel showing the same transgenic reporter line injected with a morpholino against *irf8* revealing the absence of microglia whilst OPCs remain in position. Scale bar: 20 μm. **d)** Brightfield images of neutral red stained control and *irf8* morphant animals showing the absence of phagocytic cells at 4 d.p.f. (n=21/22 animals in control/*irf8* MO, two-tailed Mann-Whitney test). **e)** Confocal images showing RGC axons in tectal neuropil. Arrows indicate ectopic branches extending outside the tectal neuropil in wildtypes, *olig2* and *irf8* morphants, respectively. Scale bar: 20 μm. **f)** Example images of single RGC axon arbors wildtypes, *olig2* morphants and *irf8* morphants. Scale bar: 10 μm.

**Supplementary Figure 4:**
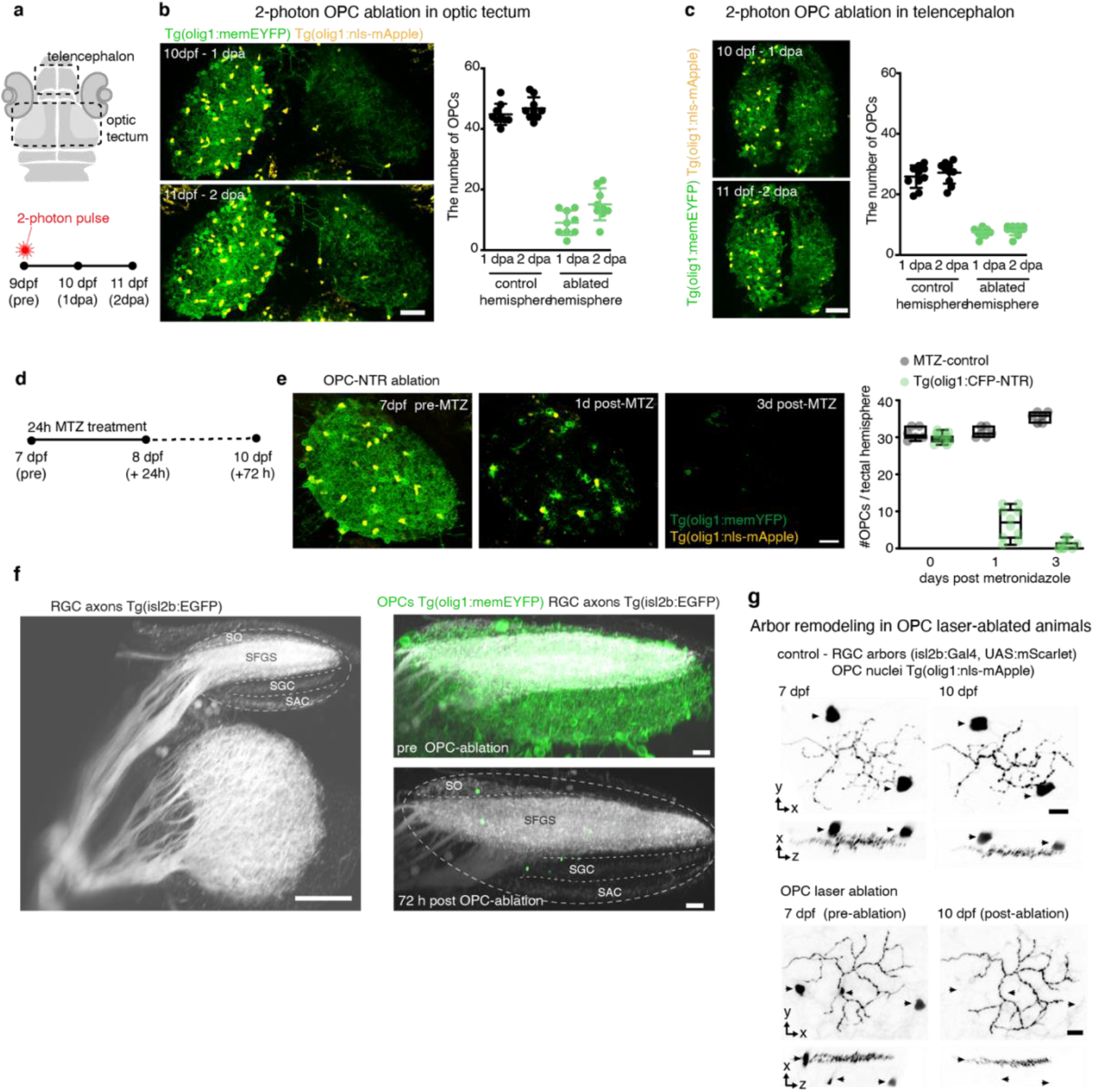
Characterization of late OPC ablation methods. **a)** Top: drawing of zebrafish head delineating the position of the optic tectum and telencephalon as shown in panels (b) and (d). Bottom: Schematic of OPC laser ablation paradigm. **b)** Dorsal views of OPC reporter lines showing the optical tectum after unilateral OPC ablation using 2 photon laser pulses and quantification of OPC numbers per tectal hemisphere (n=9 animals). Scale bar: 50 μm. **c)** Dorsal views of OPC reporter lines showing the telencephalon after unilateral OPC ablation using 2 photon laser pulses and quantification of OPC numbers per tectal hemisphere (n=10 animals). Scale bar: 50 μm. **d)** Schematic of OPC NTR ablation paradigm in panels e and f. **e)** Example images with OPC NTR ablation and quantification of OPC numbers at different time points after MTZ treatment of Tg(olig1:CFP-NTR) animals; n=8/14 animals in control/OPC NTR ablation. **f)** Left: 3D rotation view of RGC axon projections showing major projection layers in the tectal neuropil: stratum opticum (SO), stratum fibrosum et griseum superficiale (SFGS), stratum griseum centrale (SGC), stratum album centrale (SAC). Scale bar: 50 μm. Right: Overall layering of RGC projections remains intact after OPC ablation. Scale bar: 10 μm. **g)** Confocal images of individual RGC axon arbors at 7 and 10 dpf in control and OPC ablated animals. Arrowheads depict OPCs. Scale bars: 10μm. See Figure 3c for traces.

**Supplementary Figure 5:**
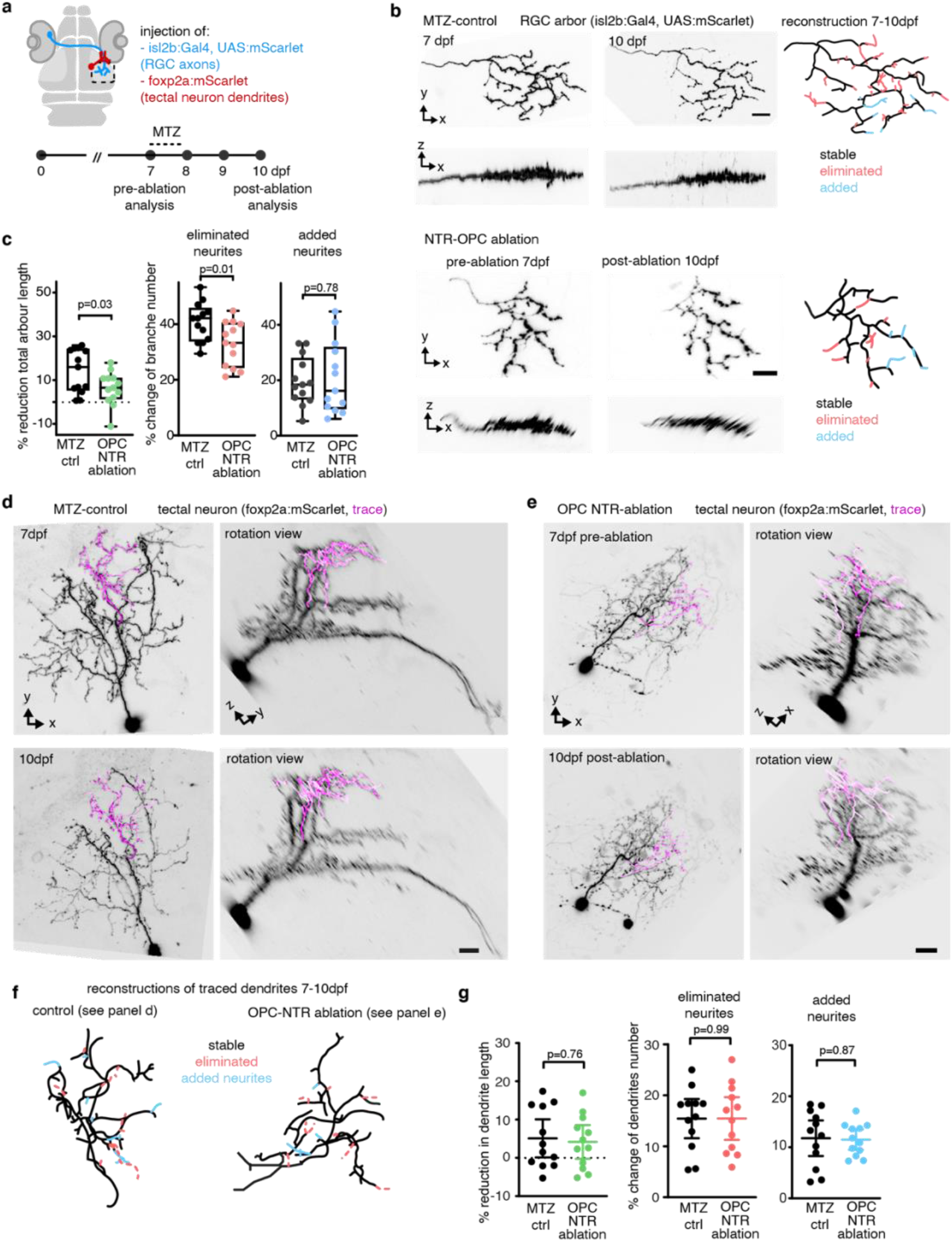
Remodeling of RGC axon arbors and periventricular interneuron (PVIN) dendrites after late OPC NTR ablation. **a)** Schematic of experimental paradigm used in this figure. **b)** Confocal images of individual RGC axon arbor in control and OPC NTR ablation animals. Tracing reveals stable, added, and eliminated processes. Scale bar: 10μm. **c)** Quantification showing RGC arbor length reduction, eliminated and added processes; n=13 cells per group in 12/10 animals in control/OPC NTR ablation (unpaired two-tailed t-test). **d** - **f)** Confocal images and rotation views of individual PVIN at two timepoints in control animals (d) and in OPC ablated animals (e). Traces show dendritic processes analyzed in panel f. Scale bars: 10μm. **g)** Quantifications of dendritic PVIN arbor remodeling between 7-10dpf in control and OPC NTR ablation animals showing no significant changes in total dendritic length, eliminated and added processes number; n=12 cells in 12 animals in control and n=12 cells in 10 animals in OPC NTR (unpaired two-tailed t test).

**Supplementary Figure 6:**
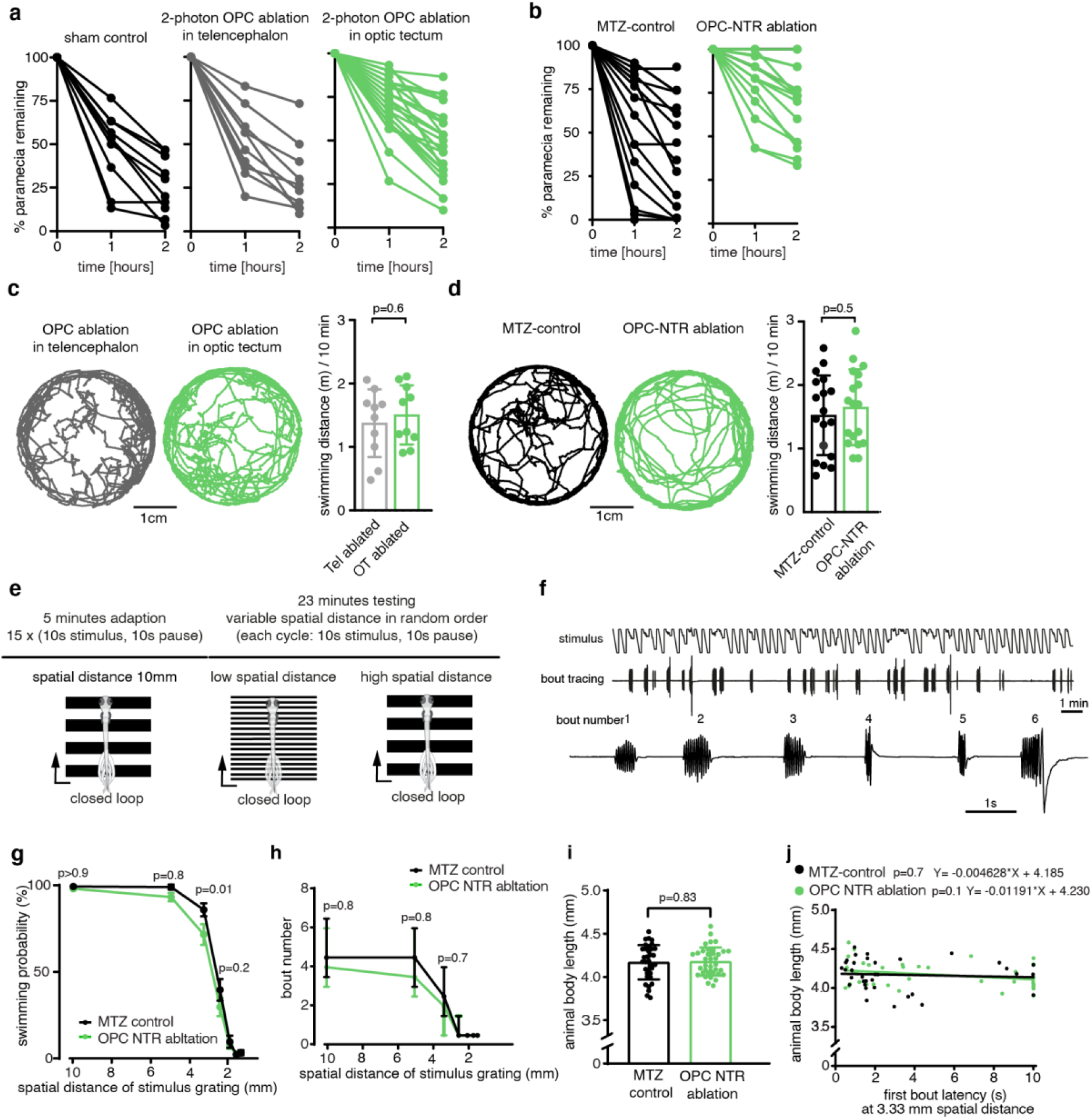
Analysis of zebrafish larvae after late OPC ablations. **a** and **b)** Quantification of paramaecium capture rates after OPC laser ablation (a) and OPC NTR ablation (b) showing measurements of individual animals over time. See also Fig. 3d and e. **c** and **d)** Example traces of the distance covered by individual freely swimming zebrafish larva within 10 minutes after OPC laser ablation (c) and OPC NTR ablation (c). **e)** Schematic of closed-loop OMR assay used. **f)** Representative tracing of an individual animal revealing different swim bouts to closed-loop stimuli in panel e. **g)** Reduced swimming probability in OPC NTR ablated animals with decreasing spatial distance of OMR stimulus grating; n=35/37 animals in control/OPC NTR (mean with SEM, two-way ANOVA). **h)** No significant changes observed in swim bout number exerted in either condition; n=35/37 animals in control/OPC NTR (median with I.Q.R., two-way ANOVA). **i)** Quantification of animal body length between control and OPC NTR animals by 3 days post MTZ treatment shows no difference between treatment groups (n=35/37 animals in control/OPC NTR, unpaired two-tailed t test). **j)** Quantification showing no correlation delay in first bout latency and animal body length. n=35/37 animals in control/OPC NTR (simple linear regression analysis).

**Supplementary Movie 1:** Animated z-projection of double transgenic olig1:memEYFP, mbp:memRFP zebrafish at 5dpf shown in Supplementary Fig. 1a.

**Supplementary Movie 2:** Animated z-projection of double transgenic isl2b:EGFP, mbp:memRFP zebrafish at 5dpf shown in Supplementary Fig. 1b.

**Supplementary Movie 3:** Animated rotation showing confocal images and tracing of individual tectal OPCs as in Supplementary Fig. 1g.

**Supplementary Movie 4:** Time-lapse of single RGC axon arbor in transgenic animals labeling all OPCs and microglia showing continuous dynamic interaction between axon and OPC processes.

**Supplementary Movie 5:** Testing optomotor response (OMR) in head-fixed larval zebrafish using Stytra software. Tail bouts are elicited by presentation of moving gratings. Tracking of initiated bouts are fed back to decrease the moving stimulus speed (closed loop).

